# SKN-1/Nrf2 regulation of neuromuscular function in response to oxidative stress requires EGL-15/FGF Receptor and DAF-2/insulin Receptor signaling in *Caenorhabditis elegans*

**DOI:** 10.1101/471755

**Authors:** Sungjin Kim, Derek Sieburth

## Abstract

The transcription factor Nrf2 plays a critical role in the organism wide-regulation of the antioxidant stress response. The Nrf2 homolog SKN-1 functions in the intestine cell non-autonomously to negatively regulate neuromuscular (NMJ) function in *Caenorhabditis elegans*. To identify additional molecules that mediate SKN-1 signaling to the NMJ, we performed a candidate screen for suppressors of aldicarb-resistance caused by acute treatment with the SKN-1 activator, arsenite. We identified two receptor tyrosine kinases, EGL-15 (fibroblast growth factor receptor, FGFR) and DAF-2 (insulin-like peptide receptor, IR) that are required for NMJ regulation in response to stress. Through double mutant analysis, we found that EGL-15 functions downstream of SKN-1 and SPHK-1 (sphingosine kinase), and that the EGL-15 ligand EGL-17 FGF and canonical EGL-15 effectors are required for oxidative stress-mediated regulation of NMJ function. DAF-2 also functions downstream of SKN-1, independently of DAF-16/FOXO, to regulate NMJ function.
Through tissue-specific rescue experiments, we found that FGFR signaling functions primarily in the hypodermis, whereas IR signaling is required in multiple tissues. Our results support the idea that the regulation of NMJ function by SKN-1 occurs via a complex organism-wide signaling network involving RTK signaling in multiple tissues.

## Introduction

The transcription factor Nrf2 plays a crucial role in the maintenance of cellular redox homeostasis by directing the expression of a cascade of antioxidant, anti-inflammatory and detoxification enzymes in response to oxidative stress (Blackwell et al., 2015). In multicellular organisms, Nrf2 activation can confer organism-wide protection from oxidative stress by regulating stress responses in distal tissues through inter-tissue signaling. In *C. elegans*, activation of the Nrf2 homolog SKN-1 in the intestine regulates longevity, survival against xenobiotics and pathogens, and neurotransmission (Kim and Jin, 2015). Although the cell-autonomous effects of Nrf2/SKN-1 in promoting cellular survival are well understood, less is known about how Nrf2/SKN-1 activation leads to changes in oxidative stress responses in distal tissues.

Oxidative stress activates Nrf2/SKN-1 by relieving it from proteolytic degradation in the cytoplasm and allowing entry into the nucleus where it regulates gene expression. Nrf2/SKN-1 activity is tightly regulated by phosphorylation and degradation (Inoue et al., 2005; Leung et al., 2014). Studies in *C. elegans* have shown that in combination with CUL-4/DDB-1, the ubiquitin ligase WD40 repeat protein WDR-23 negatively regulates the function of SKN-1 (Choe et al., 2009;

Leung et al., 2014). WDR-23 likely dissociates from SKN-1 under conditions in which oxidative stress is increased, allowing SKN-1 to accumulate in the nucleus (Choe et al., 2009). SKN-1 activation is regulated by a conserved MAP kinase cascade composed of NSY-1/MKKK, SEK-1/MKK and PMK-1/MAP kinase, which functions in the intestine to phosphorylate SKN-1 leading to its stabilization and nuclear translocation. Once activated, SKN-1 directs the expression of hundreds of genes, including genes that comprise antioxidant responses such as GST-4, glutathione-S-transferase.

We previously found that SKN-1 activation in the intestine negatively regulates NMJ function by reducing neuropeptide secretion from motor neurons. Selective activation of SKN-1 by PMK-1 or by deletion of WDR-23 in the intestine regulates NMJ function via an inter-tissue signaling mechanism, that involves the downregulation of sphingosine-1-phosphate production by intestinal SPHK-1/sphingosine kinase (Kim and Sieburth, 2018b; Staab et al., 2013). Here, we identify two pathways that function downstream or in parallel of SKN-1 to promote stress-induced regulation of NMJ function: the fibroblast growth factor receptor (FGFR) pathway, and the DAF-2 insulin receptor pathway.

FGF signaling has well defined functions in animal development, tissue repair and remodeling. In neurons, FGF signaling regulates synapse formation and refinement during development. In *C. elegans*, FGF signaling functions in several developmental processes including cell migration, muscle differentiation, axon guidance, as well as a post-developmental role in regulating fluid homeostasis (Bulow et al., 2004; Diaz-Balzac et al., 2015; Huang and Stern, 2004; Lo et al., 2008; Szewczyk and Jacobson, 2003). The FGF receptor, EGL-15 is activated by one of two FGFs, EGL-17 and LET-756. EGL-15 activates a downstream signaling cascade composed of the SEM-5/GRB2 adaptor protein, the SOS-1/guanine nucleotide exchange factor, the SOC-2/SHOC2 adaptor protein and SOC-1, a putative adaptor with a conserved PHD domain that interacts with SEM-5 (Schutzman et al., 2001).

DAF-2 is a subfamily of receptor tyrosine kinase, ortholog of insulin/IGF-1 transmembrane receptor (IR) playing a key regulator of lifespan, stress resistance, metabolism and development (Gottlieb and Ruvkun, 1994; Hung et al., 2014; Libina et al., 2003). Neuronal functions of DAF-2 include motor activity, isothermal tracking, development of cholinergic axon and touch receptor neuron (Duhon and Johnson, 1995; Hsu et al., 2009; Li et al., 2016; Murakami et al., 2005). DAF-2 also plays a role in the long term and short term learning memory (Kauffman et al., 2010).

In this study, we found that the FGF pathway consisting of EGL-17, EGL-15, SOS-1, SOC-1 and SOC-2 functions downstream of SKN-1 and SPHK-1 to negatively regulate NMJ function in response to oxidative stress. Surprisingly, EGL-15 functions primarily in the skin to regulate NMJ function. We found that DAF-2 signaling functions downstream of SKN-1 independently of DAF-16 to regulate NMJ function. Finally, we showed that DAF-2 may function in multiple distinct tissues to regulate NMJ function. These results suggest that RTK signaling in multiple tissues regulates NMJ function in response to oxidative stress.

## Material and methods

### *C. elegans* strains

All strains used in this study were maintained at 22°C following standard methods. For temperature sensitive mutants, we transferred L4 stage worms, which were grown on 22°C, to 25°C for 24 hours prior to assay. Young adult hermaphrodites were used for all experiments. The following mutant strains were used. Some of which were provided by the Caenorhabditis Genetics Center, which is funded by NIH Office of Research Infrastructure Programs (P40 OD010440): *egl-15(n1477ts), egl-15(n484ts), egl-17(ay6), let-756(s2163), sos-1(cs41ts), soc-1(n1789), soc-2(n1774), sem-5(n1799), daf-2(e1370ts), daf-2(m596ts), daf-16(mu86),* ZM8561[*daf-2(m596);hpEx2906(Pmyo-2::RFP + Prgef-1::daf-2)*], ZM8562 [*daf 2(m596);hpEx2369(Pmyo-2::RFP + Pges-1::daf-2)*], ZM8988 [*daf-2(m596);hpEx2908(Pmyo-2::RFP + Pdpy-30::daf-2)*], ZM9028 [*daf-2(m596);hpEx2905(Pmyo-2::RFP + Pmyo-3::daf-2)*], NH2447 [*ayIs2(egl-15p::GFP + dpy-20(+)*]. The *wdr-23(tm1718)* strain was provided by National BioResource Project (Japan). The wild type reference strain was N2 Bristol. The genes and mutant strains tested in our screen are listed in the Supplemental file 1.

### Molecular biology

All genes were cloned from *C. elegans* cDNA or genomic DNA from wild type worms and inserted into pPD49.26 using standard molecular biology techniques. Promoter DNA fragments were amplified from mixed stage genomic DNA. The following plasmids were generated and used: *pSK46[Pges-1::egl-15(genomic)], pSK47[Prab-3::egl-15(genomic)], pSK48[Pcol-12::egl-15(genomic)], pSK57[Pcol-12::daf-2(genomic)], pSK80[Pegl-15::gfp]*.

### Transgenic lines

Transgenic strains were generated by injecting expression constructs (10–25 ng/μl) and the coinjection marker KP#708 (*Pttx-3::rfp*, 40 ng/μl) or KP#1106 (*Pmyo-2::gfp* 10 ng/μl) into N2 or corresponding mutants. Microinjection was performed following standard techniques as previously described (Mello et al., 1991). At least three lines for each transgene were tested and a representative transgene was used for the further experiments. The following transgenic arrays were made: *vjEx1309[Pcol-12::daf-2], vjEx1239[Prab-3::egl-15], vjEx1241[Pges-1::egl-15], vjEx1243[Pcol-12::egl-15], vjEx1593[Pegl-15::gfp]*.

### Microscopy and analysis

Fluorescence microscopy experiments were performed following previous methods (Kim and Sieburth, 2018a). Briefly, for all fluorescence microscopy analysis, L4 stage or young adult worms were immobilized by using 2,3-butanedione monoxime (BDM, 30 mg/mL; Sigma) in M9 buffer then mounted on 2% agarose pads for imaging. To image and quantify the intestinal fluorescence intensity of P*gst-4::GFP and* P*egl-15::GFP* posterior intestinal cells were selected as a representative region. Images were captured with the Nikon eclipse 90i microscope equipped with a Nikon PlanApo 40 x or 60x or 100x objective (NA = 1.4) and a PhotometricsCoolsnap ES2 or a Hamamatsu Orca Flash LT+ CMOS camera. Metamorph 7.0 software (Universal Imaging/Molecular Devices) was used to capture serial image stacks, and the maximum intensity was measured (Kim and Sieburth, 2018a). Intensity quantification analysis was performed on the same day to equalize the absolute fluorescence levels between samples within same experimental set.

### RNA Interference

Feeding RNAi knockdown assay was performed following the established protocol (Kamath and Ahringer, 2003). Briefly, gravid adult animals were placed on RNAi plates seeded with HT115(DE3) bacteria transformed with L4440 vector containing fragment of knockdown genes or empty L4440 vector as a control to collect eggs then removed after 4 hours to obtain age-matched synchronized worm population. Young adult animals were used for every RNAi assay.

### Pharmacology

For aldicarb assays, the percentage of paralyzed young adult animals was counted every 10 to 15 minutes starting about one hour after placing worms on 1mM aldicarb (Bayer) plates. NGM Plates containing aldicarb were freshly made one day before each assay. Wild type animals were included in each set of assays to control for assay-to-assay variability arising from slightly different aldicarb concentrations in each batch of assay plates. Two to three replicates of at least 20 worms were performed per strain analyzed. For arsenite exposure, at least 50 young adult animals were transferred to NGM plates supplemented with 5mM sodium arsenite (RICCA Chemicals) for 4 hours prior to aldicarb assay and P*egl-15::gfp* imaging. To induce arsenite-activated P*gst-4::gfp*, at least 50 young adult animals were incubated with 5mM arsenite in M9 buffer for 1 hour, then images were taken after 4 hours of recovery.

### Statistical Analysis

For the Figure 1D and E, the Student’s t test (two-tailed) was used to determine the statistical significance. “ns” above the bars denotes P values greater than 0.05. Error bars in the figures indicate ±SEM. The numbers of animals tested are indicated in each figure.

**Figure 1.**
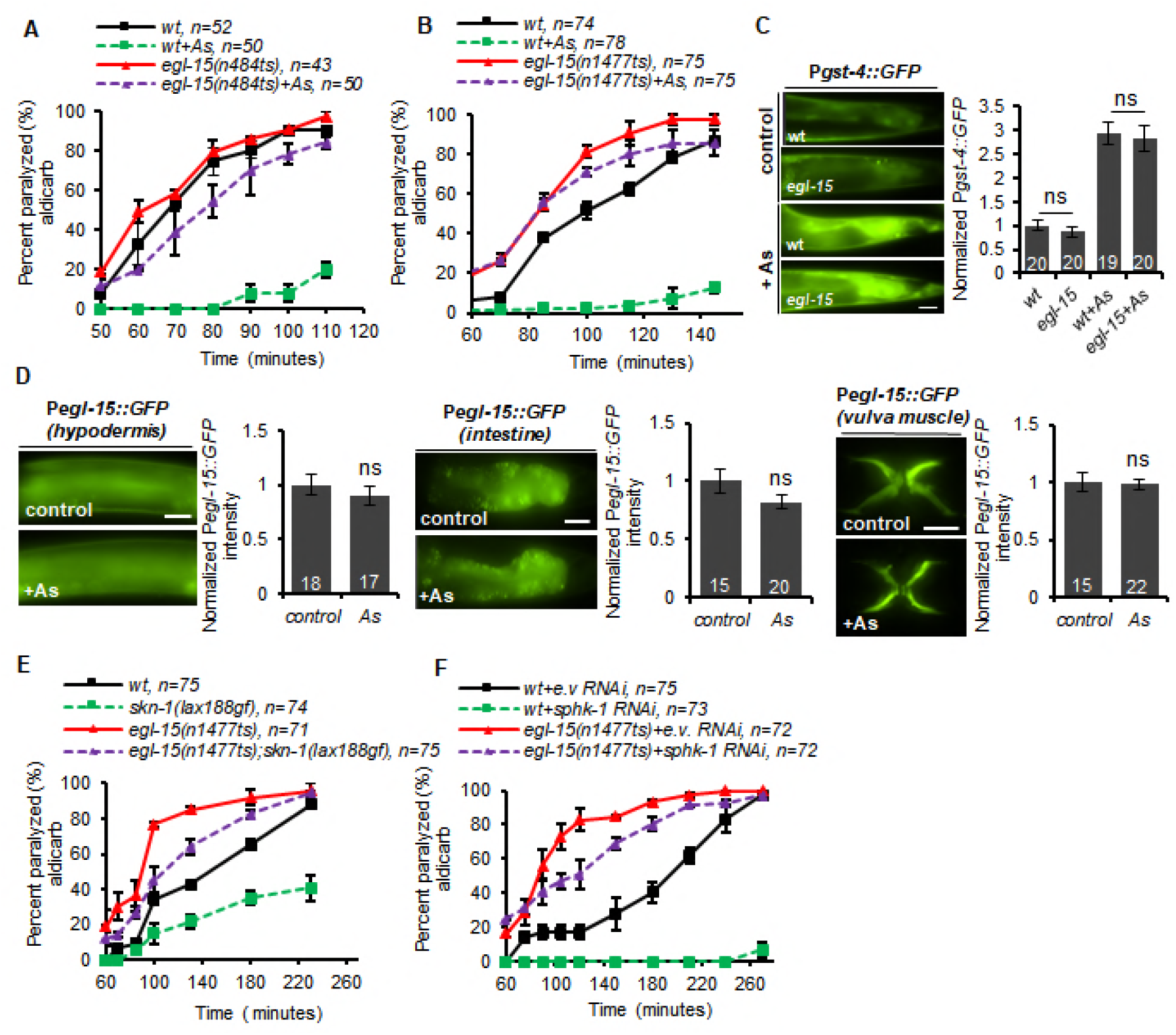
EGL-15 functions downstream of SKN-1 and SPHK-1 to promote arsenite-induced aldicarb resistance. (A-B) Time course of aldicarb-induced paralysis of wild type (wt) or *egl-15 (n1477ts) or egl-15 (n484ts)* mutants following arsenite treatment (As). (C) Representative images of posterior intestines of indicated strains expressing GFP driven under the *gst-4* promoter (*left*) in the absence or presence of arsenite (As). Quantification of the average GFP intensity (*right*). (D) Representative images and quantification of GFP driven under *egl-15* promoter in hypodermis (*upper left*), intestine (*upper right)* and vulva muscle (lower) following control or arsenite treatment. (E) Time course of aldicarb-induced paralysis of wild type control, *skn-1(lax188 gain-of-function)*, *egl-15(n1477ts)* or *skn-1(lax188gf);egl-15* double mutants. (F) Time course of aldicarb-induced paralysis of wild type (wt) and *egl-15* mutants treated with empty vector (e.v.) control or *sphk-1* RNAi. “ns” above the bars denotes P values greater than 0.05. Number of animals tested are indicated. Error bars indicate ±SEM.

### Data Availability

Strains and plasmids used in this study are available upon request. The authors clarify that all data necessary for confirming the conclusions of the findings are present within the article, figures, and supplemental file.

## Results

### A screen for genes that promote aldicarb resistance in response to oxidative stress

We previously showed that acute (4 hour) exposure to the oxidative stressor arsenite causes resistance to the paralytic effects of aldicarb (Kim and Sieburth, 2018b; Staab et al., 2013). Aldicarb is an acetylcholine esterase inhibitor, and aldicarb treatment leads to acetylcholine accumulation in synaptic clefs at neuromuscular junctions and subsequent paralysis due to muscle hyper-contraction. Animals defective in acetylcholine secretion exhibit delayed paralysis (referred to here as aldicarb resistance) because of a delay in acetylcholine accumulation in synaptic clefts. Arsenite causes aldicarb resistance by activating the SKN-1 pathway in the intestine (Kim and Sieburth, 2018b; Staab et al., 2013). To identify additional genes that mediate the effects of arsenite on neuromuscular function, we performed a large-scale candidate screen for mutants that blocked or attenuated aldicarb resistance caused by arsenite treatment. We selected 90 candidate genes based on their known roles in presynaptic function, signaling transduction, protein secretion, or stress response (Supplemental file 1). Nearly all of the mutants corresponding to these genes had wild type responses to arsenite, with the exception of *sphk-1*, *pmk-1, sek-1 and nsy-1* mutants, which were included as positive controls, as well as two additional mutants: *egl-15/*FGFR and *daf-2/*IGFR. Both *egl-*15 and *daf-2* mutants significantly attenuated the ability of arsenite treatment to cause aldicarb resistance (Supplemental file 1), revealing a potential role for FGF and IR signaling in regulating stress-induced aldicarb response.

### EGL-15 promotes arsenite-induced aldicarb resistance

*egl-15* encodes the sole *C. elegans* ortholog of the fibroblast growth factor receptor (FGFR). Temperature sensitive *egl-15* mutants are viable at the permissive temperature (22°C) and display a scrawny phenotype when grown at the restrictive temperature of 25°C (Dixon et al., 2006). *egl-15(n484ts)* mutants, which encode a W167Stop nonsense mutation, were shifted to 25°C as L4s, when development is complete, and assayed for aldicarb responsiveness as adults, 24 hours later. We found that the shifted *egl-15(n484ts)* mutants exhibited similar aldicarb sensitivity as wild type controls in the absence of arsenite. However, *egl-15(n484ts)* mutants became significantly less aldicarb resistant than wild type controls following arsenite treatment (Figure 1A). A second temperature sensitive *egl-15* mutant, *(n1477ts)*, which encodes a W930Stop nonsense mutation (DeVore et al., 1995), exhibited slight but significant hypersensitivity to aldicarb following temperature up-shift compared to wild type controls in the absence of arsenite (Figure 1B). In the presence of arsenite, *egl-15(n1477ts)* mutants remained nearly as aldicarb hypersensitive as untreated mutants (Figure 1B). Thus, EGL-15 has a post-developmental role in promoting arsenite-induced aldicarb resistance.

### EGL-15 functions downstream of SKN-1 and SPHK-1 to promote arsenite-induced aldicarb resistance

To determine whether EGL-15 is a SKN-1 activator, we examined the requirement of *egl-15* in the stress-induced expression of *gst-4*/glutathione-S-transferase, which is a direct transcriptional target of SKN-1 (Choe et al., 2009; Wu et al., 2016). A transcriptional reporter in which GFP is expressed under the *gst-4* promoter (P*gst-4::GFP*) is expressed at low levels in the absence of stress whereas arsenite treatment significantly increases P*gst-4::GFP* expression in the intestine in a *skn-1*-dependnet manner (Choe et al., 2009; Wu et al., 2016). We found that the expression of P*gst-4::GFP* in the intestine was similar in *egl-15* mutants and wild type controls in the absence of arsenite (Figure 1C). Following four hour arsenite treatment, P*gst-4::GFP* expression increased about three-fold in wild type controls, and we observed a similar three-fold increase in *egl-15* mutants (Figure 1C). Thus, both baseline SKN-1 activity and arsenite-induced activation of SKN-1 are normal in *egl-15* mutants, suggesting that EGL-15 does not regulate SKN-1 activity.

To determine whether EGL-15 is a transcriptional target of SKN-1, we examined the effects of arsenite treatment on the fluorescence intensity of P*egl-15::GFP* reporters, in which the 2.0 kb promoter fragment of *egl-15* drives the expression of GFP. We found that transgenic animals expressing P*egl-15::GFP* exhibited fluorescence in the intestine, vulval muscle and hypodermis, in agreement with prior studies (Bulow et al., 2004; Mounsey et al., 2002). Arsenite treatment did not significantly alter the fluorescence intensity of P*egl-15::GFP* in any of these tissues (Figure 1D). These results suggest that EGL-15 is not transcriptionally regulated by SKN-1 activation.

To determine whether EGL-15 functions in the SKN-1 pathway to regulate aldicarb resistance, we examined *skn-1* mutants that are constitutively active. The *skn-1(lax188gf)* mutation alters an amino acid in a protein-interaction domain that renders SKN-1 constitutively active (Paek et al., 2012). *skn-1(lax188gf)* mutants are aldicarb resistant (Staab et al., 2013). We found that *egl-15* mutations significantly reduced the aldicarb resistance of *skn-1(lax188gf)* mutants (Figure 1E). SKN-1 activation leads to aldicarb resistance in part by negatively regulating SPHK-1 signaling in the intestine (Kim and Sieburth, 2018b). To determine whether EGL-15 mediates the effects of SPHK-1 in this pathway, we examined aldicarb responses of *egl-15* mutants where intestinal *sphk-1* activity is knocked down by RNAi. Animals treated with *sphk-1* RNAi are strongly resistant to aldicarb (Chan et al., 2012; Kim and Sieburth, 2018b). Strikingly, *egl-15* mutations nearly completely suppressed the aldicarb resistance caused by *sphk-1* RNAi (Figure 1F). These results indicate that EGL-15 and SPHK-1 function antagonistically to regulate aldicarb responsiveness, and that EGL-15 functions downstream of or in parallel to SPHK-1. Together, these results reveal a role for EGL-15 in regulating neuromuscular function in response to SKN-1 activation.

### The FGF ligand EGL-17 mediates arsenite-induced aldicarb resistance

Two FGF-related ligands signal through EGL-15 to control distinct functions of EGL-15. EGL-17/FGF promotes sex myoblast migration while LET-756/FGF promotes axon guidance and fluid homeostasis (Birnbaum et al., 2005; Burdine et al., 1997; DeVore et al., 1995; Diaz-Balzac et al., 2015; Lo et al., 2008; Popovici et al., 1999; Sundaram et al., 1996). To determine whether either of these FGF ligands promotes stress-induced aldicarb resistance, we examined aldicarb responses of *egl-17* and *let-756* mutants. *egl-17(ay6)* is a deletion allele that is predicted to eliminate *egl-17* activity (Burdine et al., 1997). *egl-17(ay6)* mutants displayed wild type aldicarb responsiveness in the absence of arsenite. Following arsenite treatment, *egl-17(ay6)* mutants became more aldicarb resistant than untreated mutants, but this shift in aldicarb resistance was significantly smaller than that of wild type controls treated with arsenite (Figure 2A). This partial response suggests that EGL-17 contributes to arsenite-induced aldicarb resistance, but is not likely to be solely responsible for EGL-15 activation in this process.

**Figure 2.**
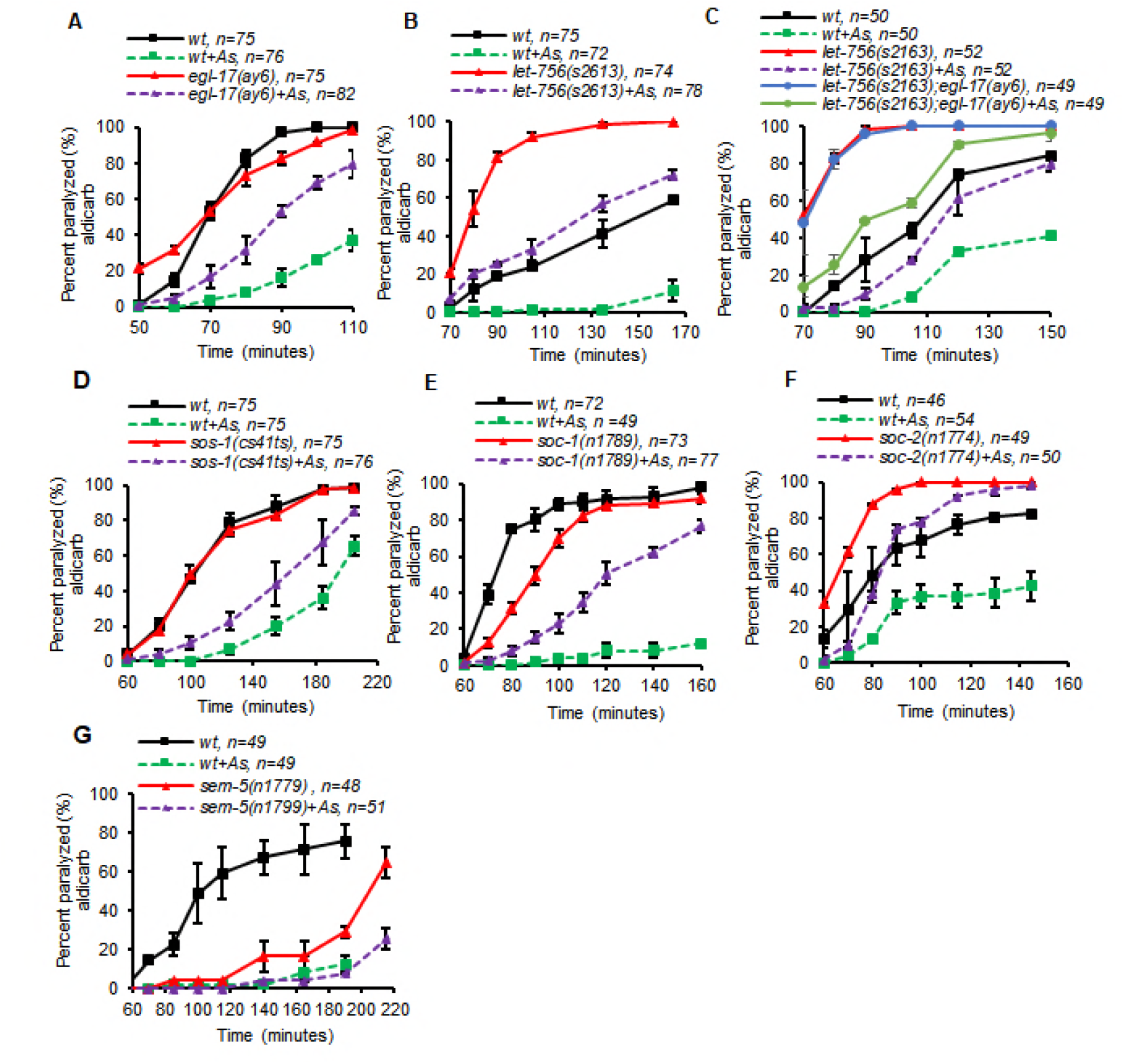
EGL-17 FGF and the SOS-1/SOC-1/SOC-2 signaling cascade is required for arsenite-induced aldicarb resistance. (A-G) Time course of aldicarb-induced paralysis of indicated strains in the absence or presence of arsenite. Number of animals tested are indicated. Error bars indicate ±SEM.

To test the role of LET-765, we examined *let-756(s2613)* hypomorphic mutants, which are viable, unlike null mutants, which die as larvae (Popovici et al., 2004). *let-756(s2613)* mutants were significantly more sensitive to aldicarb-induced paralysis than wild type controls in the absence of arsenite (Figure 2B), revealing a role for LET-765 in inhibiting NMJ function in the absence of stress. Arsenite treatment caused a large shift toward aldicarb resistance in *let-756* mutants that was similar to that of wild type controls (Figure 2B), suggesting that stress-induced aldicarb resistance is not impaired in these mutants. The partial defect in arsenite-induced aldicarb resistance of *egl-17* mutants was not enhanced by *let-756* mutations (Figure 2C). Thus, we conclude that EGL-17 contributes to arsenite-induced aldicarb resistance, and LET-756 may not. However, because the *let-756* mutant analyzed was not null, it is not possible to make a definitive determination of its contribution in this pathway.

### The SOS-1/SOC-1/SOC-2 signaling cascade mediates arsenite-induced aldicarb resistance

We next examined mutants corresponding to each of the known components that make up the signaling cascade downstream of EGL-15 for defects in arsenite-induced aldicarb resistance. *sos-1(cs41ts)* mutants are temperature sensitive (Abdus-Saboor et al., 2011), and when shifted to 25°C for 24 hours prior to assay, exhibited wild type aldicarb responsiveness in the absence of stress. *sos-1(cs41ts)* mutants became significantly less aldicarb resistant following arsenite treatment than wild type controls (Figure 2D). *soc-1(n1789)* and *soc-2 (n1774)* are null and hypomorph alleles, respectively (Schutzman et al., 2001). We found that *soc-1(n1789)* mutants were slightly aldicarb resistant in the absence of arsenite whereas *soc-2 (n1774)* mutants were hypersensitive to aldicarb (Figure 2E, F). However, both mutants became significantly less aldicarb resistant than wild type controls following arsenite treatment (Figure 2E, F). Finally, *sem-5(n1779)* hypomorphic mutants were extremely resistant to aldicarb in the absence of arsenite, revealing a role of SEM-5 promoting NMJ function. Arsenite treatment elicited a further increase in aldicarb resistance in *sem-5(n1779)* mutants, but this shift was much smaller than the shift elicited in wild type controls (Figure 2G). Thus, SOS-1, SOC-1 and SOC-2 contribute to stress-induced aldicarb resistance. The contribution of SEM-5 is more difficult to ascertain given that the mutants were so resistance to aldicarb in the absence of stress. Because EGL-15 signaling is left partially intact in each of the signaling mutants tested, we conclude that these components may function in parallel with each other, or may function with other unidentified components activated by EGL-15 to contribute to stress-induced aldicarb responsiveness.

### EGL-15 functions in the hypodermis to promote arsenite-induced aldicarb resistance

EGL-15 has been reported to be expressed in multiple tissues, including the hypodermis, vulval muscles, intestine and neurons (Bulow et al., 2004; Huang and Stern, 2004). To determine the site of action of EGL-15 with respect to stress-induced aldicarb resistance, we performed tissue-specific rescue experiments by generating transgenic *egl-15(n1477ts)* mutants expressing *egl-15* genomic DNA selectively in the hypodermis (using the *col-12* promoter (Olofsson, 2014)), intestine (using the *ges-1* promoter(Kim and Sieburth, 2018a)) or nervous system (using the *rab-3* promoter (Kim and Sieburth, 2018b)). Expression of *egl-15* in each of these tissues did not rescue the weak aldicarb hypersensitivity of *egl-15(n1477ts)* mutants in the absence of stress, suggesting that the basal response of *egl-15(n1477ts)* mutants to aldicarb may require EGL-15 signaling in multiple tissues or in other tissue(s) not tested here (Figure 3A-C). However, EGL-15 expression in the hypodermis fully restored arsenite-induced aldicarb resistance to *egl-15(n1477ts)* mutants (Figure 3A). In contrast, expression of *egl-15* transgenes in the nervous system failed to rescue the aldicarb sensitivity of *egl-15(n1477ts)* mutants (Figure 3B), and expression of *egl-15* in the intestine partially restored stress-induced aldicarb resistance to *egl-15(n1477ts)* mutants (Figure 3C). These results suggest that EGL-15 signaling in the hypodermis is critical for arsenite-induced aldicarb resistance. Our results support a model whereby under conditions of low stress, EGL-15 signaling is inhibited by SPHK-1 in the intestine. During stress, SPHK-1 activity is inhibited by SKN-1 activation, which in turn leads to increased EGL-15 signaling in the skin (and possibly also the intestine) and negative regulation of neuromuscular function (Figure 3D).

**Figure 3.**
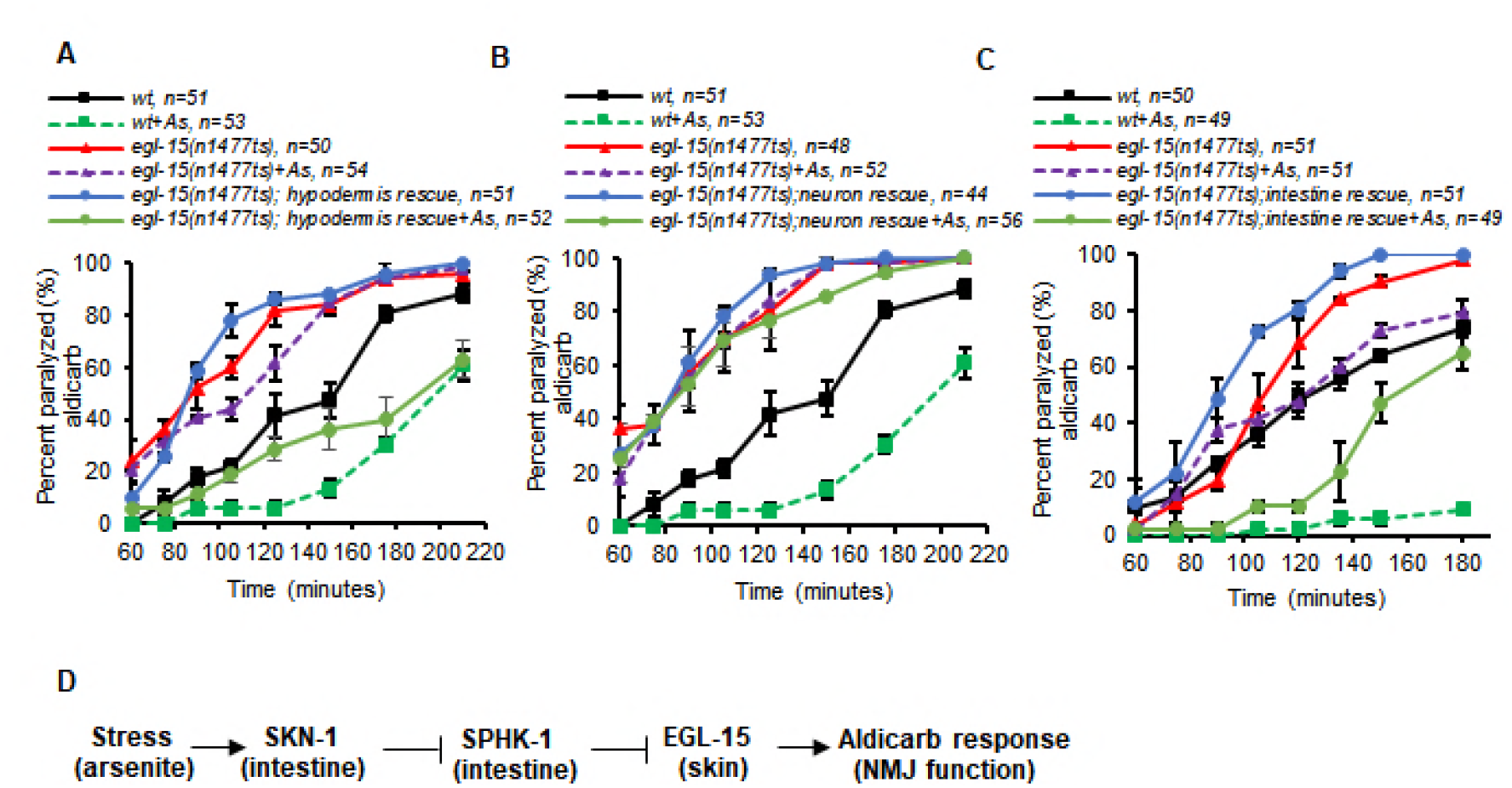
EGL-15 is required in the hypodermis to regulate arsenite mediated aldicarb resistance. (A-C) Time course of aldicarb-induced paralysis of the indicated strains in the absence or presence of arsenite (As). “hypodermis rescue, neuron rescue, intestine rescue,” denotes *egl-15* transgene expression using the tissue-specific promoters *col-12, rab-3*, or *ges-1,* respectively. (D) Schematic working model whereby EGL-15 signaling in the skin (and intestine) is negatively regulated by SPHK-1 signaling in the intestine during low stress conditions. Following SKN-1 activation by oxidative stress, SPHK-1 activity is inhibited, which in turn leads to increased EGL-15 activity resulting in aldicarb resistance. Number of animals tested are indicated. Error bars indicate ±SEM.

### DAF-2 is required for aldicarb resistance induced by oxidative stress

The second gene identified in our screen was *daf-2*. We examined two temperature sensitive *daf-2* mutants, *daf-2(e1370ts),* which encodes a P1547S missense mutation, and *daf-2(m596ts),* which encodes a G471S missense mutation (Bulger et al., 2017). Both mutants are viable at the semi-permissive temperature of 22°C and enter into the dauer stage at the restrictive temperature of 25°C (Bulger et al., 2017). In the absence of arsenite, both *daf-2(e1370ts)* and *daf-2(m596ts)* mutants were slightly more aldicarb resistant than wild type controls when cultured at 22°C (Figure 4A and 4E). Following arsenite treatment, *daf-2(e1370ts)* mutants failed to become more aldicarb resistant than untreated mutants, remaining nearly as sensitive to aldicarb as untreated *daf-2* controls (Figure 4A). These results reveal a role for DAF-2 in promoting arsenite-induced aldicarb resistance in adult animals that is distinct from its role in regulating development.

**Figure 4.**
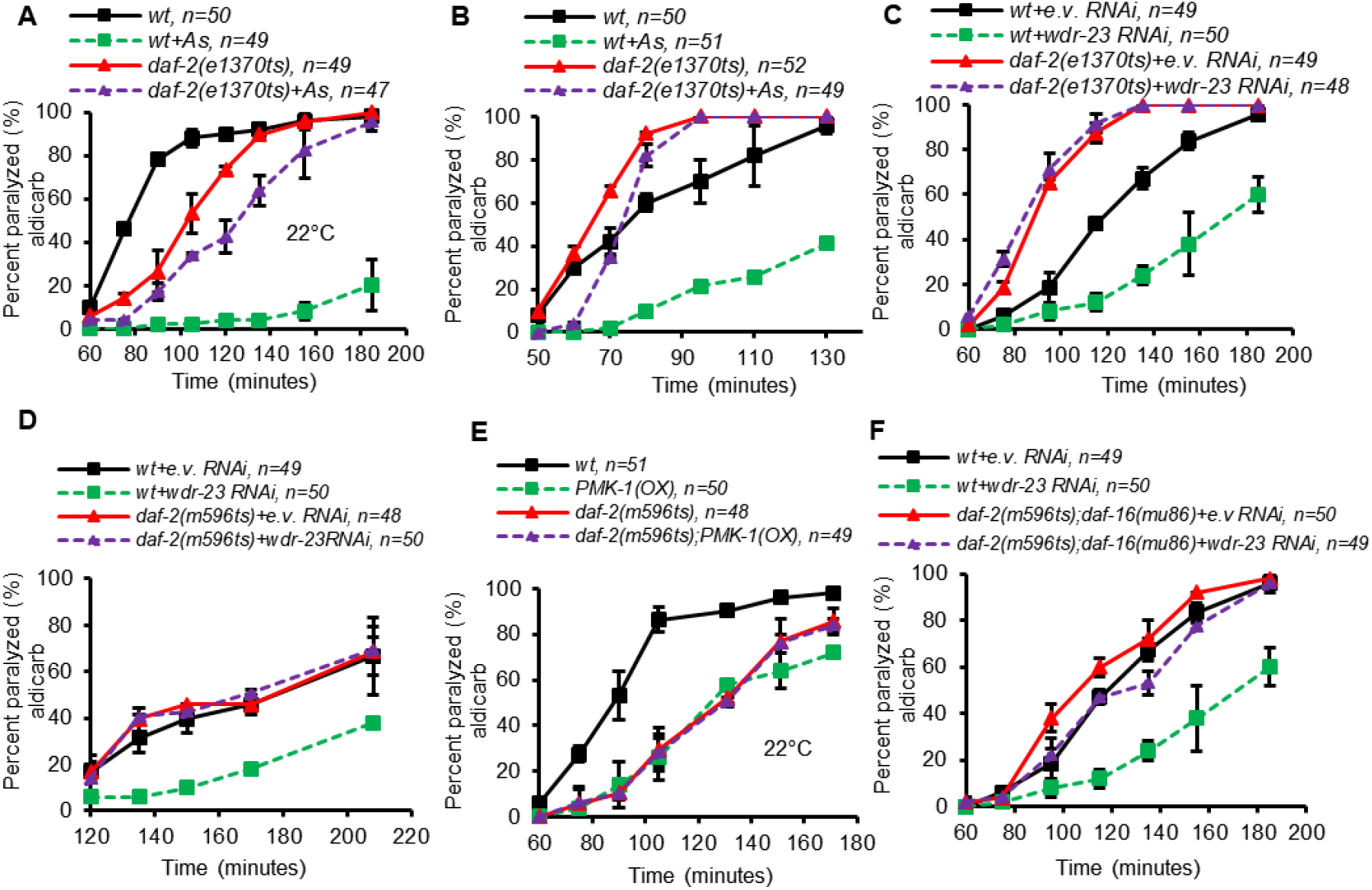
DAF-2 functions downstream of SKN-1 to regulate NMJ function. (A-B) Time course of aldicarb-induced paralysis of wild type control (wt) and *daf-2(e1370ts)* mutants grown at 22°C (A) or 25°C for 24 hours (B) prior to aldicarb assays following control or arsenite treatment. (C-D) Time course of aldicarb-induced paralysis of indicated strains treated with empty vector (e.v.) control or *wdr-23* RNAi. (E) Time course of aldicarb-induced paralysis of indicated strains. PMK-1(OX) denotes animals over-expressing *pmk-1* transgenes in the intestine. (F) Time course of aldicarb-induced paralysis of indicated strains following empty vector (e.v.) control or *wdr-23* RNAi treatment. Number of animals tested are indicated. Error bars indicate ±SEM.

### DAF-2 functions downstream or in parallel to SKN-1

To determine whether DAF-2 functions in the SKN-1 pathway to regulate aldicarb responsiveness, we examined the aldicarb responsiveness of *daf-2* mutants in which SKN-1 is constitutively active. WDR-23 promotes neuromuscular function by negatively regulating SKN-1 in the intestine. *wdr-23* null mutants show delayed aldicarb paralysis in the absence of stress that is completely dependent upon *skn-1* and is not enhanced by arsenite treatment (Kim and Sieburth, 2018b; Staab et al., 2013). As expected, *wdr-23* knockdown by RNAi led to strong aldicarb resistance. However, knockdown of *wdr-23* was unable to cause aldicarb resistance in either *daf-2(e1370ts)* or *daf-2(m596ts)* mutants (Figure 4C, D), suggesting that the aldicarb resistance phenotype caused by SKN-1 activation is dependent upon *daf-2* signaling. PMK-1 positively regulates SKN-1 activity in the intestine by phosphorylating SKN-1, leading to its stabilization and its accumulation in the nucleus (Inoue et al., 2005). Animals over-expressing PMK-1 cDNA specifically in intestine (under the *ges-1* promoter) exhibited enhanced aldicarb resistance compared to non-transgenic controls (Figure 4E and (Kim and Sieburth, 2018b)).

However, PMK-1 overexpression was unable to make *daf-2(m596ts)* mutants more aldicarb resistant than non-transgenic controls (Figure 4E). Together, these results are consistent with a function of DAF-2 either downstream or in parallel to SKN-1 in regulating aldicarb resistance in response to stress.

DAF-2 signaling exerts is biological effects by either negatively regulating the DAF-16/FOXO transcription factor (Chen et al., 2013; Simon et al., 2014; Sun et al., 2017)), or by a mechanism that is independent of DAF-16 (Szewczyk et al., 2007). If DAF-2 negatively regulates DAF-16 in this stress response, we predict that *daf-16* mutations should restore SKN-1-induced aldicarb resistance to *daf-2* mutants. We found that *daf-2; daf-16* double mutants exhibited aldicarb responsiveness that was similar to *daf-2* single mutants (Figure 4F). However, *daf-16* mutations did not restore aldicarb resistance to *daf-2* mutants treated with *wdr-23* RNAi (Figure 4D and 4F). This result suggests that DAF-2-mediated regulation of aldicarb resistance does not require DAF-16 signaling.

### DAF-2 functions in multiple tissues to promote arsenite-induced aldicarb resistance

DAF-2 is expressed in multiple tissues including intestine, nervous system, hypodermis and muscle (Hunt-Newbury et al., 2007; McKay et al., 2003). To identify the tissue in which DAF-2 functions in SKN-1-mediated aldicarb resistance, we performed a series of tissue-specific rescue experiments using *daf-2* genomic DNA. A prior study generated a panel of extrachromosomal arrays in which *daf-2* genomic DNA was expressed in different tissues (Hung et al., 2014). We examined strains bearing these extrachromosomal arrays for their ability to restore normal stress-induced aldicarb resistance to *daf-2(m596ts)* mutants. To activate the stress response, we knocked down *wdr-23* by RNAi in *daf-2(m596ts)* mutants. As expected, expression of *daf-2* genomic DNA in all tissues (using the *dpy-30* promoter) fully reverted aldicarb resistance to *daf-2* mutants treated with *wdr-32* RNAi (Figure 5A). Next, we tested whether DAF-2 expression selectively in the intestine (using the *ges-1* promoter), the nervous system (using the *rab-3* promoter), the hypodermis (using the *col-12* promoter), or in body wall muscle (using the *myo-3* promoter) could restore normal aldicarb responsiveness to *daf-2(m596ts)* mutants in which *wdr-23* was knocked down. Interestingly, we found that expression of DAF-2 in any single tissue failed to rescue *daf-2(m596ts)* mutants (Figure 5B-E). These results suggest that DAF-2 may function in more than one tissue to regulate SKN-1 mediated aldicarb resistance. Alternatively, DAF-2 may function in a tissue not tested here, such as the germ line to regulate aldicarb responsiveness.

**Figure 5.**
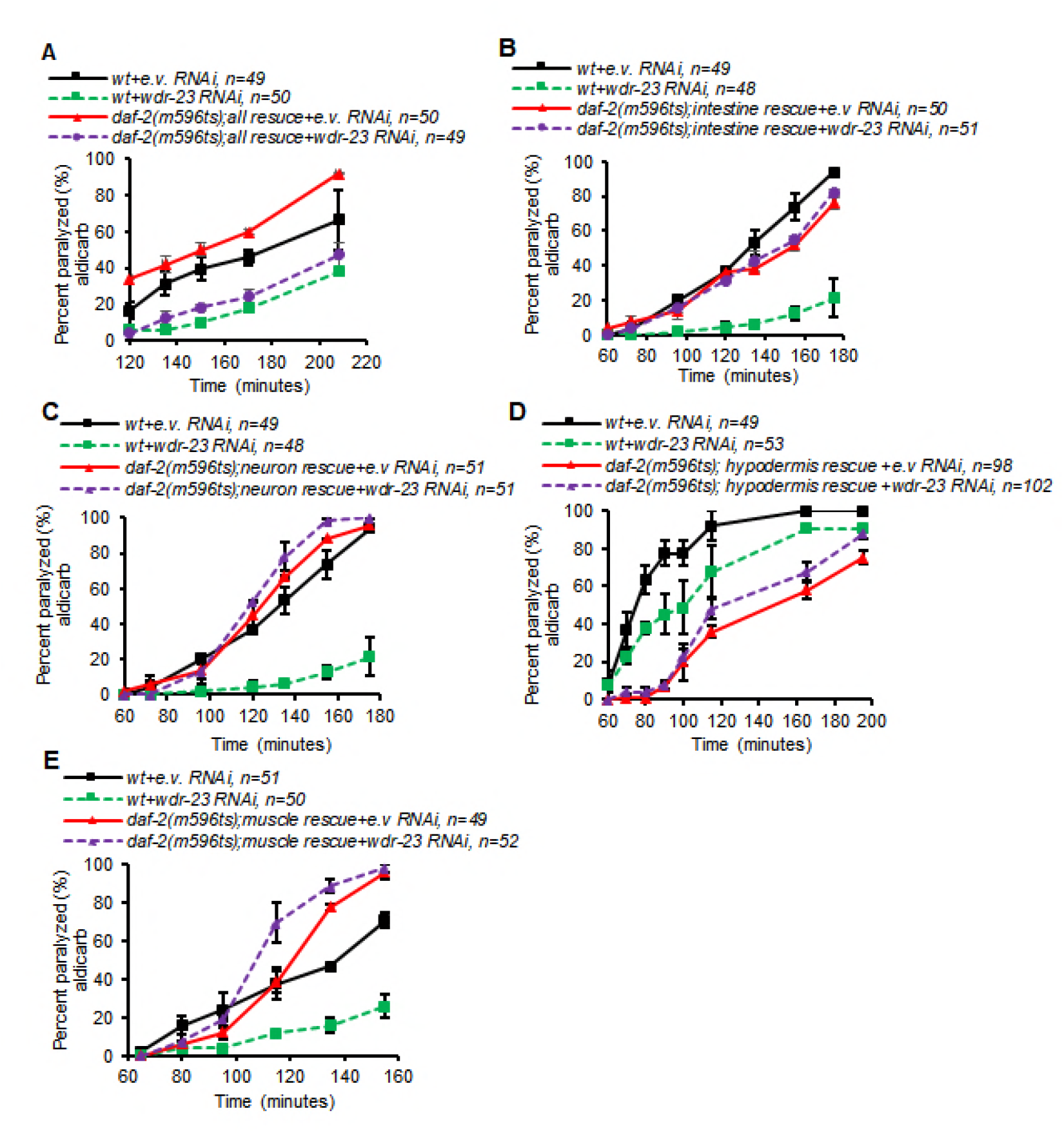
DAF-2 functions in multiple tissues to regulate SKN-1 dependent NMJ function. (A-E) Time course of aldicarb-induced paralysis of wild type (wt) animals or *daf-2(m596ts) transgenic* mutants following empty control (e.v.) or *wdr-23* RNAi treatment. “all rescue, intestine rescue, neuron rescue, hypodermis rescue, muscle rescue” denotes DAF-2 transgenes expressing under the tissue-specific promoters *dyp-30, ges-1, rab-3, col-12 or myo-3*, respectively. Number of animals tested are indicated. Error bars indicate ±SEM.

## Discussion

Multicellular organisms are exposed to variety of stresses in the form of endogenous or environmental stress-induced by changes in their surroundings. Therefore, they have developed complex organism-wide defense mechanism to prevent the cellular damage and maintain proper cellular homeostasis. Our results support the idea that the regulation of NMJ function by oxidative stress is mediated by complex signaling networks that act across multiple tissues and involve at least two RTK signaling cascades, FGFR and IR. Our genetic analysis revealed that both of these RTKs likely function downstream of SKN-1 to promote aldicarb resistance caused by acute oxidative stress. Our temperature shift experiments show that these RTK pathways regulate NMJ physiology rather than development. We found that EGL-15 functions primarily in the hypodermis to regulate NMJ function, whereas DAF-2 signaling may be required in multiple different tissues. Thus, EGL-15 and DAF-2 signaling may mediate inter-tissue communication between the intestine, where SKN-1 is activated and the NMJ during the oxidative stress response.

Aldicarb resistance can arise from defects in acetylcholine release from NMJs, neuropeptide secretion or muscle excitability. We previously found that the aldicarb resistance caused by SKN-1 activation is not due to enhanced detoxification of aldicarb or alterations in muscle excitability but instead is due to reduction in neurotransmitter release from motor neurons (Staab et al., 2013). We subsequently found that inhibition of intestinal SPHK-1 by SKN-1 results in a reduction in neuropeptide secretion from motor neurons (Kim and Sieburth, 2018b). Since EGL-15 is required for the effects of SPHK-1 on NMJ function, we speculate that FGF signaling may also regulate neuropeptide secretion, although further analysis will be needed to determine the detailed mechanism by which FGFR and IR signaling regulates NMJ function.

### FGF signaling in regulating NMJ function

Our results are consistent with the idea that EGL-17 activates EGL-15 to regulate aldicarb response. The *egl-15* gene encodes two receptor isoforms, EGL-15(A) and EGL-15(B) generated by alternative splicing, that differ in their extracellular domains and are proposed to have different ligand binding specificity. EGL-17 binds to EGL-15(A) to mediate migration of sex myoblasts(SM) while LET-756 binds to EGL-15(B) to regulate development and neuronal growth (Birnbaum et al., 2005). Because *egl-17* null mutants did not fully block arsenite induced aldicarb resistance, it is possible that EGL-17 and LET-756 or another unidentified ligand may function redundantly to regulate aldicarb responsiveness.

Our results show that mutations in any single known downstream component of the EGL-15 pathway attenuate but do not block the arsenite induced aldicarb resistance, suggesting that there may be signaling redundancy among these components, or there may be other unidentified EGL-15 effectors. In mammals, the FGFR activates multiple different cytosolic signaling factors in a signal and context dependent manner. Additional downstream targets of FGFR that were not tested here include the adapter proteins FRS2a and CRKL, and well as STAT family members (Ornitz and Itoh, 2015). It will be interesting to determine whether these genes function with SOC-1 and/or SOS-1 in this pathway. Interestingly, we found that in the absence of arsenite, *let-756* or *soc-2* mutants were significantly hypersensitive to aldicarb-induced paralysis, whereas *soc-1* and *sem-5* mutants were resistant to aldicarb, revealing previously unreported roles for these signaling components in negatively and positively regulating NMJ function, respectively. The aldicarb resistance of *sem-5* mutants complicates the interpretation of the relatively small shift to aldicarb resistance in *sem-5* mutants upon arsenite treatment: it may reflect a requirement of SEM-5 in promoting arsenite responsiveness, or it could represent a ceiling effect given the extreme aldicarb resistance of the mutant under baseline conditions. Further experiments will be needed to determine whether SEM-5 is involved in this pathway.

How does EGL-15 signaling regulate aldicarb responsiveness in response to stress? EGL-15 has a post-developmental function in regulating fluid homeostasis (Huang and Stern, 2004). However, defects in fluid homeostasis are not likely to account for the aldicarb phenotypes of *egl-15* mutants, since *let-756* mutants, which are also defective in fluid homeostasis, did not block arsenite-induced aldicarb resistance, whereas *egl-17* mutants, which are not defective in fluid homeostasis, behaved similarly to *egl-15* mutants. Thus, the aldicarb phenotypes of the FGF mutants do not correlate with defects in fluid homeostasis. EGL-17 release from the hypodermis is required for sex myoblast migration during development, and in adults, EGL-17 is expressed in several cells of the ventral in the hypodermis, as well as in a pharyngeal neuron (Burdine et al., 1998; Dixon et al., 2006; Hunt-Newbury et al., 2007). Thus, increased EGL-17 secretion from either of these sites may activate EGL-15 signaling in the skin in response to stress.

### Insulin-like receptor signaling in regulating NMJ function

Our results suggest that the insulin receptor DAF-2 regulates SKN-1 mediated NMJ function in response to oxidative stress, and that DAF-2 function is likely required in multiple tissues. Selective expression of DAF-2 in neurons and muscle restores hypoxic death to *daf-2* mutants which are highly resistant to hypoxia (Scott et al., 2002). Neuronal DAF-2 is involved in life span extension and dauer formation of *daf-2* mutants whereas DAF-2 function in muscle, which is required for fat metabolism, is not critical for life span extension and dauer phenotype (Wolkow et al., 2000) suggesting that biological function of DAF-2 is highly tissue-specific. We speculate that DAF-2 associated NMJ function in response to stress may be occurring in multiple tissues by inter-tissue signaling mediated by one or more of the 40 insulins in *C. elegans* (Zheng et al., 2018). We showed that DAF-2 functions independently of DAF-16 to regulate NMJ function. Similarly, DAF-2 functions independently of DAF-16 in muscles to inhibit protein degradation and promote proper mobility (Szewczyk et al., 2007). Notably, we found that selective expression of *daf-2* in the hypodermis led to aldicarb resistance (Figure 5D), a phenotype that was not observed when expressing *daf-2* in any other tissue tested, suggesting that enhanced DAF-2 signaling in the hypodermis may regulate NMJ function in the absence of stress. Indeed, DAF-2 signaling has been implicated in decreasing motor function in aged animals, but its site of action was not determined (Liu et al., 2013).

Our results suggest that DAF-2 signaling functions downstream of SKN-1 to regulate NMJ function. Consistent with this, tyrosine phosphorylation of the insulin receptor substrates 1 and 2 (IRS-1 and −2) by the insulin receptor is strongly reduced in the Nrf2-deficient mice (Beyer and Werner, 2008). Furthermore, insulin receptor tyrosine kinase is greatly activated by oxidative stressor hydrogen peroxide (Droge, 2005). Interestingly, DAF-2 signaling has been implicated in the activation of SKN-1 during aging through the regulation of the Akt kinases AKT-1/2 (Tullet et al., 2008). The function of DAF-2 downstream of SKN-1 that we report here is likely to be distinct from the function identified for DAF-2 during ageing, further underscoring the complex role of insulin signaling in the SKN-1 pathway.

## Acknowledgements

We thank members of the lab for advice and critical review of the manuscript, and Mei Zhen for providing DAF-2 plasmids. This work was supported by grants from the NIH National Institute of Neurological Disorders and Stroke (NINDS) to D.S. (NS071085 and NS099414). Some strains were provided by the Caenorhabditis Genetics Center (CGC), which is funded by the NIH Office of Research Infrastructure Programs (P40 OD010440).

